# Long term consumption of green tea EGCG enhances healthspan and lifespan in mice by mitigating multiple aspects of cellular senescence in mitotic and post-mitotic tissues, gut dysbiosis and immunosenescence

**DOI:** 10.1101/2021.01.01.425058

**Authors:** Rohit Sharma, Ravi Kumar, Anamika Sharma, Abhishek Goel, Yogendra Padwad

**Author notes:** Address correspondence to: Dr. Rohit Sharma, INSPIRE Faculty Fellow, CSIR-Institute of Himalayan Bioresource Technology, Palampur, India-176061. ORCID iD: 0000-0001-6209-3845., Dr Yogendra Padwad, Senior Scientist, CSIR-Institute of Himalayan Bioresource Technology, Palampur, India-176061.

## Abstract

Cellular senescence is emerging as the causal nexus of aging, and its potential modulators present an effective strategy to counter age-related morbidity. The current study profiled the extent of cellular senescence in different organs of mice at four different time-points of lifespan, and explored the influence of epigallocatechin gallate (EGCG) consumption in impacting multiple aspects of aging biology. We report that adipose and intestinal tissues are highly vulnerable to cellular senescence as evident by age-associated increase in DNA damage response, activation of cell cycle inhibitors (p53/p21) and induction of SASP (p38MAPK/NF-κB/Cox-2). Further, a distinct modulation of nutrient signaling pathway mediators (AMPK/Akt/SIRT3 and 5), and a decrease in autophagy effectors was also observed in aging animals. Systemic inflamm-aging markers (TNF-α/IL-1β) and splenic CD4/CD8 ratio increased with age, while NK cell population decreased. Metagenomic analyses revealed age-related decrease in the diversity of microbial species while an increase in the abundance of various pathogenic bacterial genera was also observed. Long term EGCG consumption enhanced lifespan of animals by attenuating markers of DNA damage, cell cycle inhibitors and SASP in adipose, intestine and liver tissue. Mechanistically, EGCG inhibited the activation of AMPK and Akt and enhanced mitochondrial SIRT3 and SIRT5 expression, as well as autophagic response in adipose and intestinal tissues. Systemic presence of inflamm-aging markers decreased while expression of T cell immune response regulator CD69 increased in EGCG fed animals. EGCG also improved age-related decrease in the diversity of microbial species and suppressed the growth of pathogenic microbes. In short, our results provide compelling evidence that post-mitotic adipose tissue is a major site of cellular senescence and SASP activation, and that chronic EGCG consumption can influence several aspects of aging and senescence resulting in improved organismal healthspan and lifespan.

## 1. Introduction

Aging represents a unique risk factor that predisposes elderly to inflammatory disorders and infectious diseases ultimately increasing the rate of morbidity and mortality. This is of particular concern since the global elderly population is steadily rising, and is projected to more than double in the year 2050 (UN report, 2019). Recent attempts to understand the underlying molecular mechanisms of aging have emphasized a central role of the process of cellular senescence (van Deursen, 2014). It is rapidly emerging that age-associated gradual accumulation of SC and chronic presence of SASP creates pro-tumorigenic and pro-inflammatory environment which is detrimental to tissue/organ functions (Prattichizzo et al 2016). Although key questions addressing the extent of cellular senescence, particularly in the post-mitotic tissues such as adipose (Sapieha and Mallette, 2018), or those relating cellular senescence and immune dysfunctions are least understood (Sharma and Padwad, 2020a); yet several emerging reports suggest that selective ablation of senescent cells through senolytics and/or suppressing the development of cellular senescence/SASP can have strong implications in promoting longevity and attenuating the susceptibility to age-related disorders (Baker et al., 2011,2016; Palmer et al., 2019). In addition, it is also increasingly recognized that the accumulation of oxi-inflammatory stress in cells as well as functional inefficacy of the immune system play a critical role in the development and progression of cellular senescence and its known effects during aging (Song et al 2020). Thus, it is evident that strategies aimed to regress (or at least delay) the different aspects of cellular senescence and/or immunosenescence are highly desirable in the quest for improving lifespan and attenuating age-associated morbidity and mortality.

Nutritional intervention during aging (nutrigerontology) is considered a pragmatic approach for enhancing the quality of life in elderly (Verburgh, 2015). This is perhaps most strikingly observed in the case of plant polyphenols which have been correlated with improved indices of morbidity and mortality both in experimental animals as well as in epidemiological studies (Sharma and Padwad, 2020b; Wu et al 2020). EGCG is the most abundant polyphenol in green tea and is associated with anti-inflammatory, anti-cancerous and anti-obesity attributes. It has also been observed that EGCG consumption can augment lifespan in rats (Niu et al 2013) as well as improve healthspan in worms (Xiong et al 2018). However, studies delineating the role of EGCG in influencing the different facets of cellular senescence vis-à-vis aging are rare and also limited to in vitro analyses only (Han et al 2012; Kumar et al 2019). Considering these lacunae, the present in vivo study was designed to address two fundamental questions: First, we aimed to assess how cellular senescence progresses in different animal organs with an emphasis on the post-mitotic adipose tissue. Second, we sought to understand the influence of EGCG consumption in animal physiology through a holistic assessment of various markers of cellular senescence, immunosenescence and gut dysbiosis during aging. Our results show that adipose tissue is highly vulnerable to oxi-inflammatory stress and cellular senescence, and that EGCG consumption can effectively suppress several indicators of senescence thereby promoting organismal healthspan and lifespan.

## 2. Materials and methods

### 2.1 Animal experimental design

Male swiss albino mice (2 months old) were procured from the animal house facility of CSIR-IHBT, Palampur. This particular species and sex of animals were chosen based on our previous study (Sharma et al 2017). Animals were divided into eight groups of 12 each: 4 control groups and 4 EGCG treated groups. All animals were maintained on commercial diet pellets (Golden Feeds, New Delhi, India) and housed in the experimental animal facility of CSIR-IHBT, Palampur under standard laboratory conditions (12:12 h light/dark cycle, 22 ±2°C temperature, 50-60% humidity). EGCG was orally administered at a concentration of 100 mg/kg/animal body weight by dissolving in drinking water based on our previous observation (Sharma et al 2017). To ensure proper consumption and adequate distribution, EGCG was always freshly prepared in distilled water and served to animals during morning hours (9:00 AM Indian Standard Time) in standard water bottles at 5 ml per animal per day equaling the approximate daily liquid consumption of each mice. No additional water was subsequently provided to the animals. Animals were appropriately sensitized to this feeding regimen before beginning of actual experiments. All control animals followed the same protocol, but without EGCG. A total of four different time points in mice lifespan viz. 6,10,14 and18 months were chosen to profile age-associated changes as well as the effects of EGCG. For this, after every four months of respective dose regimen, at least six animals from both control and EGCG groups each were sacrificed and blood, peritoneal fluid, small intestine, subcutaneous adipose tissue, liver and spleen of the animals were isolated for downstream processing and analyses. Body weight and food consumption of animals were regularly monitored, and periodic examination by a veterinarian was conducted for signs of infection or disease. All animal experiments were performed as per the guidelines and approval of the institutional animal ethics committee (Approval no. IAEC/IHBTP-1/May 2018, dated May 10, 2018).

### 2.2 Isolation of peritoneal macrophages

Peritoneal macrophages were isolated as described previously (Sharma et al 2018). Briefly, the mice peritoneal cavity was flushed with Dulbeco’s Modified Eagle Medium (DMEM) Ham’s F12 (without phenol red), and after gentle massaging, the media was carefully recollected from the cavity avoiding any tissue. The isolated cell suspension was then incubated overnight in a humidified 5% CO_2_ incubator at 37°C to allow attachment of adherent cells. Subsequent washing removed any non-adherent cells, and the adherent macrophages were used to isolate RNA and perform qRT-PCR.

### 2.3 Splenocytes isolation and immunophenotyping

Splenocytes were isolated and used for immunophenotyping as described previously (Sharma et al 2018). FITC-labelled anti-mouse CD3e, APC labelled anti-mouse CD4, PE-labelled anti-mouse CD8, PE-labelled anti-mouse NKp46 (CD335), and vioblue-labelled anti-mouse CD69 (Miltenyi Biotec, Germany) were used for identification of cell sub-populations and activation status. NK cells were identified as CD3-NKp46+ (Joyce and Pollard, 2009; Mair et al 2016), and the expression of CD69 was considered as marker of early T cell activation and regulation. Briefly, isolated splenocytes were incubated with respective antibodies for 10 min at 4°C, followed by washing and resuspension of the pelleted cells in FACS buffer for data acquisition. Cells were subjected to flow cytometry using AMNIS ImageStream®X Mark II Imaging Flow Cytometer, and data were analyzed using the INSPIRE ImageStream system software. Lymphocyte population was gated using forward and side scatter profiles after identifying single circular cells using the area versus aspect ratio feature. Fluorescence minus one control were used in all experiments to precisely identify the specific sub-population (Mahnke and Roederer, 2007).

### 2.4 Cytokines estimation

Systemic inflammation in plasma was assessed by measuring cytokine levels using commercially available quantitative sandwich ELISA kits for interleukin 1 beta (IL-1β) (Invitrogen, USA, Cat no.:88-7013-88), and tumor necrosis factor α (TNF-α) (eBioscience, USA, Cat no.: 88-7324-88) in undiluted samples as per manufacturer’s protocol.

### 2.5 Total RNA isolation and qRT-PCR

Total cellular RNA was isolated by using commercially available Qiagen RNeasy mini kit (74104, Qiagen, Germany). RNA was quantified and qualitatively assessed using a nanodrop instrument following which one step qRT-PCR was performed using QuantiFast SYBR Green PCR kit (204054; Qiagen, Germany). Table 1 lists the primers used for mRNA quantification and cellular glyceraldehyde-3-phosphate dehydrogenase expression was used as a normalizing control. PCR reactions were performed using the Step one Plus™ Real-Time PCR system (Applied Biosystems, USA), and ΔΔCt method was used to determine the relative mRNA quantification (Sharma et al. 2017).

**Table 1.**
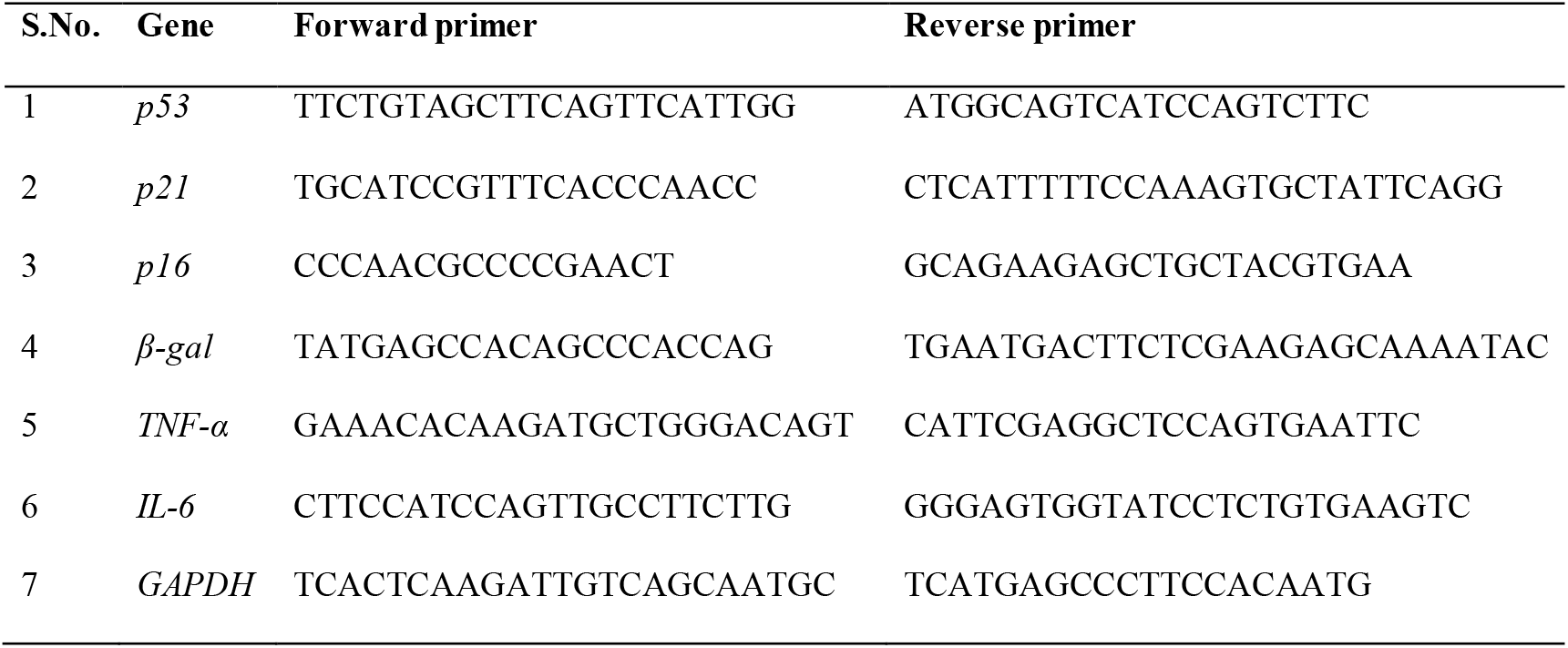
List of primer sequences used for qRT-PCR.

### 2.6 Protein isolation and western blotting

A section of each tissue, viz. adipose, intestine and liver were collected and snap frozen at −80°C until use. For isolation of total proteins, each tissue section was first washed twice with PBS (10 mM) and then lysed in RIPA buffer (R0278; Sigma Aldrich, USA). After centrifugation of lysates, protein quantity in the supernatant was determined by Bradford assay (Bradford, 1976). For western blotting, total protein (50 μg) was subjected to SDS-PAGE followed by transfer to polyvinylidene difluoride membrane (GE Healthcare Life Sciences, Europe) using a semidry trans-blotter (BioRad, USA). Non-fat dried milk (5%) was used for subsequent blocking of the membranes at room temperature and after washing, the membranes were probed with the following primary antibodies overnight at 4°C with gentle shaking: anti-p53 (13-4100; 1:500) and anti-p21^WAF1^ (AHZ0422; 1:250) antibodies were purchased from Thermo Fisher Scientific, USA while anti-ATM/ATR substrate (2851; 1:500), anti-p38MAPK (9212S; 1:1000), anti-p-p38MAPK (4511S; 1:1000), anti-NF-κB (8242S; 1:1000), anti-p-NF-κB (3031S; 1:500), anti-Cox2 (12282S; 1:1000), anti-AMPKα (2603; 1:1000), anti-p-AMPKα (2535S; 1:1000), anti-Akt (9272; 1:500), anti-SIRT3 (5490S; 1:1000), anti-p-Akt (9271S; 1:250), anti-SIRT5 (8782S; 1:1000); anti-Atg5 (12994; 1:1000) and anti-LC3A/B (12741; 1:1000) antibodies were procured from Cell Signaling Technology, USA. HRP-conjugated secondary antibody (7074S; Cell Signaling Technology, USA) was used for further incubation of membranes at room temperature for 1 h. After washing, protein bands were detected using Clarity Western ECL Substrate (BioRad, USA). Protein expression was quantified using GelAnalyzer software (version 19.1), and target protein band intensities were normalized using the Ponceau stain for total loaded protein (Smith et al 2018).

### 2.7 Microbial DNA extraction and whole genome metagenomic sequencing

Fecal samples were collected at the end of each respective feeding regimen and stored at −80°C till further analysis. The QIAGEN DNA Stool Mini-Kit (QIAGEN, Hilden, Germany) was used for extracting DNA from fecal samples as per manufacturer’s protocol. DNA integrity and quantity were checked using agarose gel electrophoresis and nanodrop analysis. Whole genome metagenomic sequencing (ion torrent platform with single end base pair chemistry), taxonomic annotation and analyses were outsourced from Neuberg Center of Genomic Medicine, Neuberg Supratech Micropath Laboratory & Research Institute Pvt. Ltd, Ahmedabad, Gujarat, India. Operational taxonomic units (OTUs) were identified and used for taxonomic assignment and were mapped with the 16S green genes database. The resulting taxonomy assigned OTU were then used to ascertain the abundance of microorganisms at each level.

### 2.8 Statistical analyses

Data were analyzed by using GraphPad Prism software. Results are expressed as means ± SD. Differences between the means were tested for statistical significance using a 2-way analysis of variance (ANOVA) followed by Sidak post hoc test. The significance level was set at 5% (*P* < 0.05) for all calculations.

## 3. Results

### 3.1 EGCG consumption improves animal survival

As shown in Figure 1A, the Kaplan –Meier survival curve of the control and EGCG groups began to diverge at 32 weeks and remained separated. The Mantel–Cox log rank test showed significant differences between the survival curves of EGCG and control groups, and Logrank based hazard ratio analyses indicated an average 46.13% lower risk of death in EGCG fed animals. The feed consumption in all animals started declining after 56 weeks (Figure 1B), while body weight increased steadily up to 48 weeks before showing a decline (Figure 1C). No statistical difference in the body weight or feed consumption between groups could be observed suggesting that the apparent increase in the lifespan of EGCG treated animals may be independent of changes in body weight or food intake.

**Figure 1.**
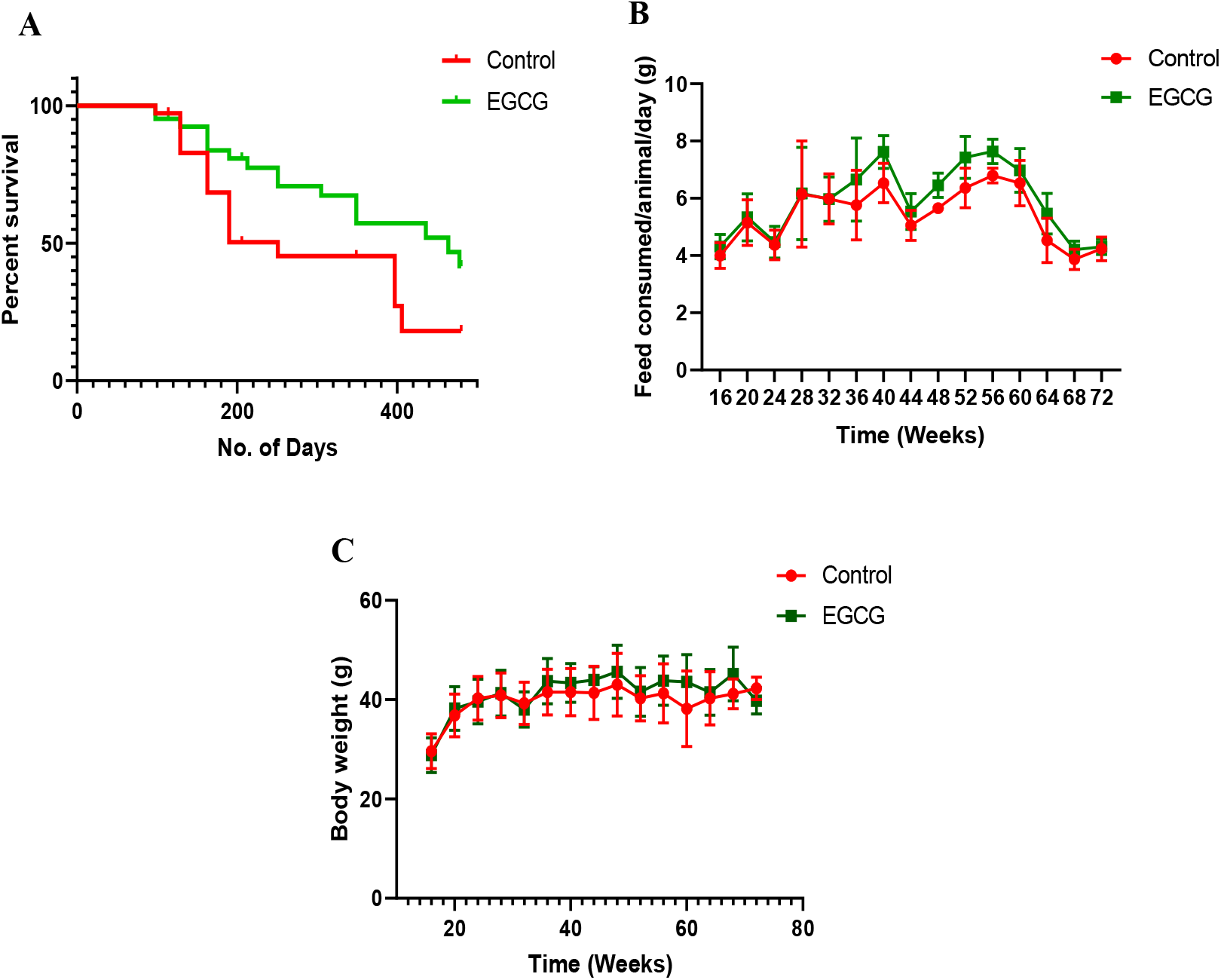
Effect of EGCG on lifespan, feed consumption and body weight in healthy mice. Animals were divided into 4 control groups and 4 EGCG fed groups, and one group each from control and EGCG groups was sacrificed after every 4 months of feeding till 18 months of animal age. **(A)** Kaplan-Meier survival curves. Hazard ratio for EGCG was 0.538 **(B)** Feed consumption **(C)** Body weight. Data are presented as means ± SD and *P* < 0.05 was considered significant.

### 3.2 Rate of cell senescence is tissue dependent which is mitigated by EGCG administration

To understand the onset and extent of cellular senescence in vivo, we evaluated markers of both senescence and SASP in the adipose tissue, intestine and liver. As shown in Figure 2A-C, protein levels of cell cycle inhibitors p53 and p21 steadily increased in the adipose tissue of control animals from 14 months of age as compared to younger animals. On the other hand, EGCG administered group showed a marked and significant decrease in both p53 and p21 expression especially in 18 months old animals (Figure 2A-C). Similarly, in the intestine tissue, a significant increase in p53 expression and a non-significant increase in p21 expression was observed in 18 months old animals which were effectively suppressed in EGCG group (Figure 2D-F). Liver also appeared to show increased p53 expression with age, while no such trend was apparent in the levels of p21 (Figure 3G-I). Nonetheless, EGCG treatment decreased p53 and p21 expression in liver suggesting robust influence of EGCG on cell cycle progression across various tested tissues (Figure 3G-I). Except for liver, a significant presence of multiple phosphorylated ATM/ATR substrates in 14-18 months old animals was also observed confirming widespread age-related DNA damage as compared to younger animals which appeared to be attenuated in EGCG fed animals (Figure 2C,F,I).

**Figure 2.**
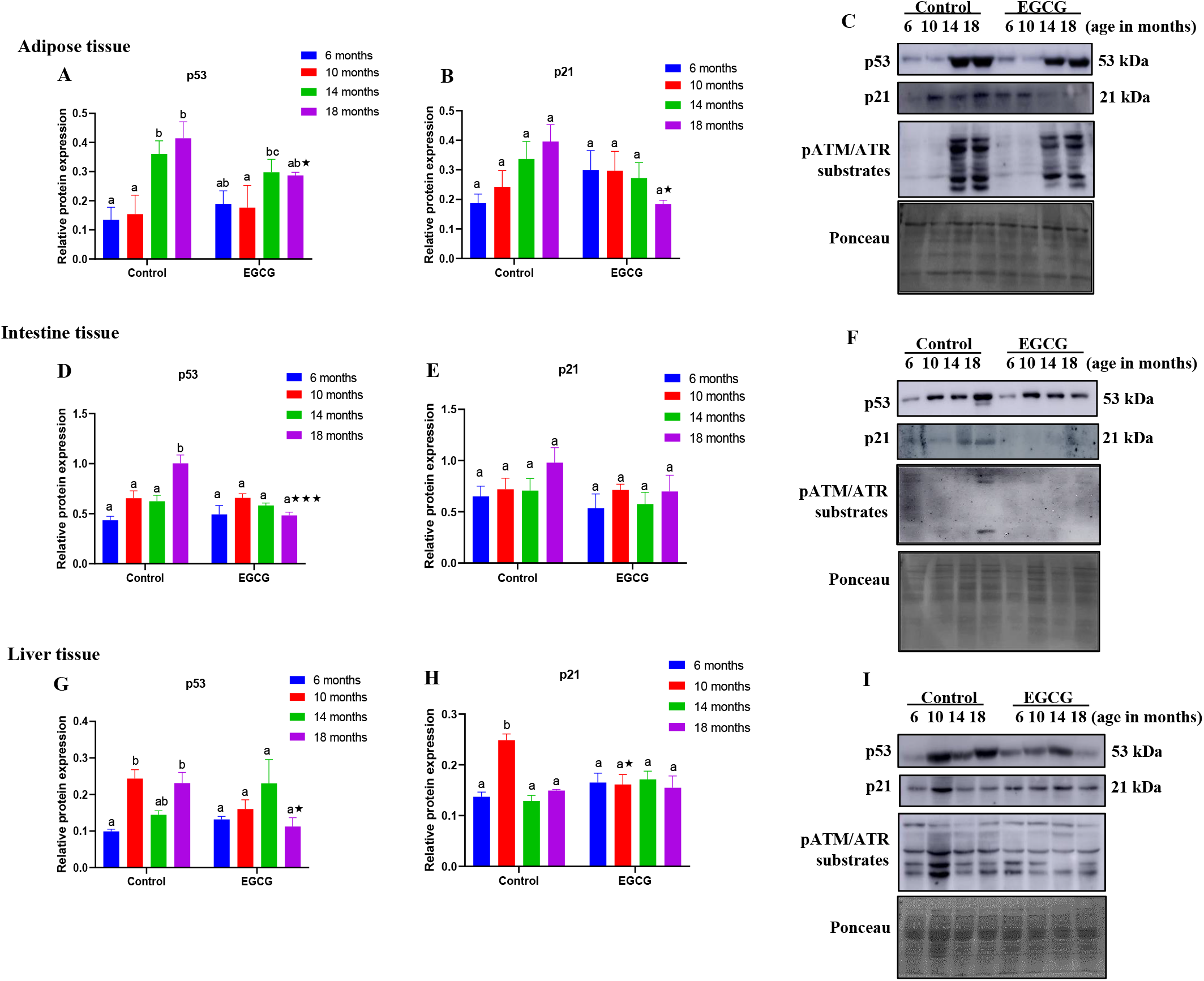
Influence of EGCG consumption on age-associated activation of cell cycle inhibitors in various tissues. Animals were divided into 4 control groups and 4 EGCG fed groups, and one group each from control and EGCG groups was sacrificed after every 4 months of feeding till 18 months of animal age. Relative protein expression in adipose tissue **(A)** p53 **(B)** p21 **(C)** Representative western blot images. Relative protein expression in intestine tissue **(D)** p53 **(E)** p21 **(F)** Representative western blot images. Relative protein expression in liver tissue **(G)** p53 **(H)** p21 **(I)** Representative western blot images. Data are presented as means ± SD (n=3) and tested using a 2-way ANOVA followed by Sidak post hoc test. Means that do not share a common letter within a group indicate statistical difference at *P* < 0.05. *represents statistical difference between control and EGCG treated groups at *P* < 0.05.

**Figure 3.**
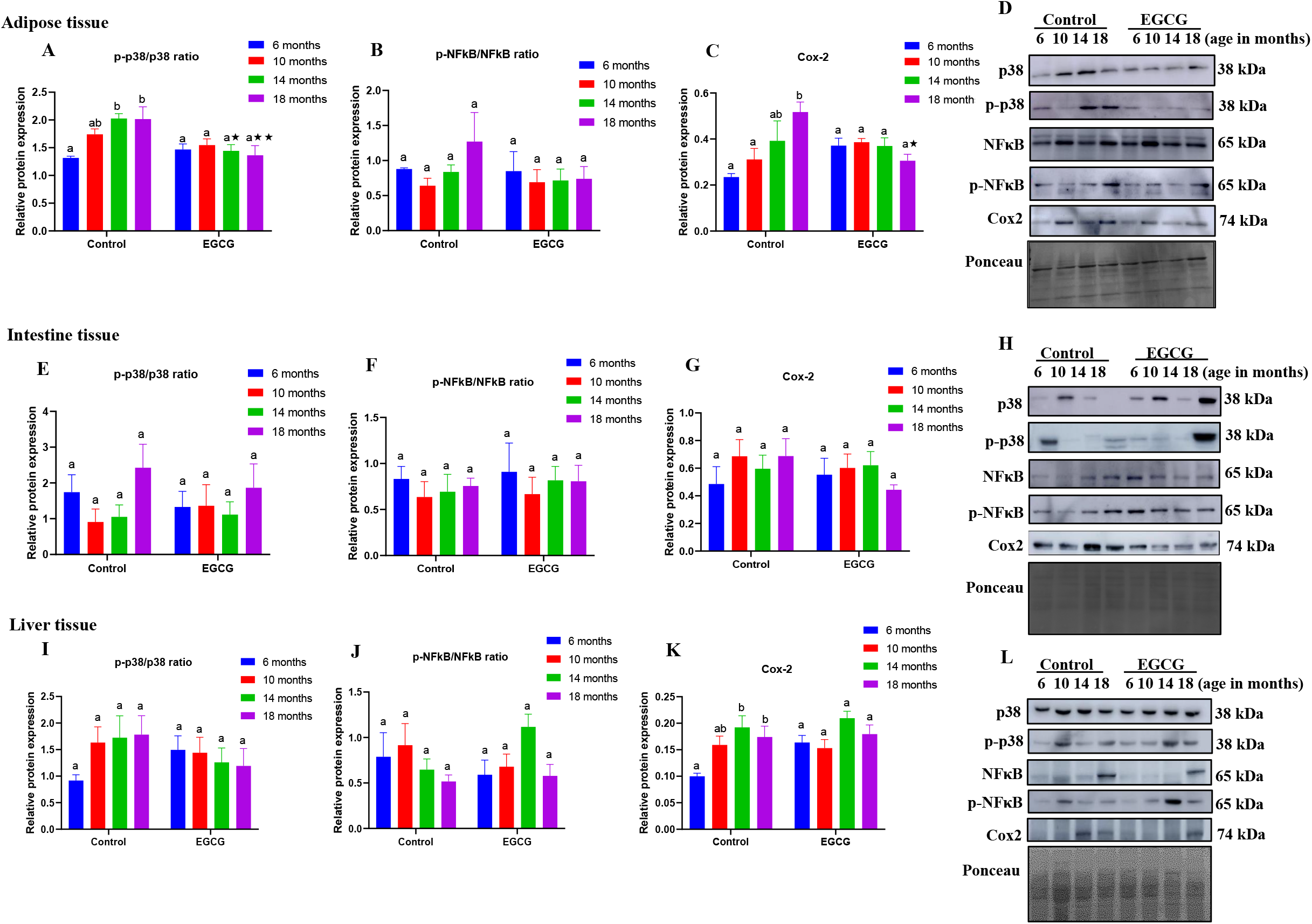
Effect of EGCG consumption on markers of SASP activation in various tissues. Animals were divided into 4 control groups and 4 EGCG fed groups, and one group each from control and EGCG groups was sacrificed after every 4 months of feeding till 18 months of animal age. Relative protein expression in adipose tissue **(A)** p-p38/p38 ratio **(B)** p-NF-κB/NF-κB ratio **(C)** Cox-2 **(D)** Representative western blot images. Relative protein expression in intestine tissue **(E)** p-p38/p38 ratio **(F)** p-NF-κB/NF-κB ratio **(G)** Cox-2 **(H)** Representative western blot images. Relative protein expression in liver tissue **(I)** p-p38/p38 ratio **(J)** p-NF-κB/NF-κB ratio **(K)** Cox-2 **(L)** Representative western blot images. Data are presented as means ± SD (n=3) and tested using a 2-way ANOVA followed by Sidak post hoc test. Means that do not share a common letter within a group indicate statistical difference at *P* < 0.05. *represents statistical difference between control and EGCG treated groups at *P* < 0.05.

### 3.3 Markers of SASP are most distinct in adipose tissue which are attenuated by EGCG administration

A stable cell senescence is characterized by a strong signature of pro-inflammatory transcription factors and effector molecules characteristic of SASP. To assess this, we evaluated protein expression and activation status of inflammatory transcription factors p38MAPK and NF-κB as well as cox-2 in various tissues. As observed with cell cycle inhibitors and DNA damage response, the adipose tissue exhibited a remarkable age-associated increase in the activation of p38MAPK and expression of cox-2 in 14-18 months old animals as compared to younger animals while a non-significant increase in NF-κB expression was also evident (Figure 3A-D). On the other hand, EGCG fed animals showed a significant suppression in the activation of p38MAPK and cox-2 in 14-18 months old animals, while a non-significant decrease in NF-κB phosphorylation was observed (Figure 3A-D). In the intestine, a gradual yet non-significant increase in the activation of p38MAPK and cox-2 was evident in 18 months old animals as compared to younger animals of the control group (Figure 3E-H). On the other hand, a slight decrease in the expression of these pro-inflammatory markers was recorded in the EGCG treated group animals (Figure 3E-H). However, liver did not reveal any specific trend although p-p38MAPK and Cox-2 expression appeared to increase with age, which were non-significantly attenuated by EGCG administration (Figure 3I-L). As a result, the adipose and intestinal tissues were utilized for further analyses since they showed robust age-associated activation of cellular senescence and SASP.

### 3.4 Age-associated deregulation in nutrient sensing pathways is influenced by EGCG

Nutrient sensing pathways are critical regulators of cell metabolism, stress response as well as senescence. In the adipose tissue, it was observed that the presence of activated AMPK and Akt were gradually increasing with age as compared to younger animals which, however, appeared to be significantly downregulated on treatment with EGCG (Figure 4A-B). On the other hand, expression of mitochondrial sirtuins SIRT3 and SIRT5 initially increased with age, but was significantly downregulated in 18 months old animals (Figure 4C-D). However, EGCG treated animals showed a general tendency of a non-significant increase in the expression of both SIRT3 and SIRT5 with age (Figure 4C-D). Representative western blot images are presented in Figure 4E. A remarkably similar pattern was also observed in the intestinal tissue wherein an age-associated gradual and significant increase in AMPK and Akt activation was accompanied by an initial increase in SIRT3 and SIRT5 expression which eventually dramatically decreased in 18 months old animals (Figure 4F-I). Conversely, EGCG treatment in 18 months old animals significantly suppressed age-associated increase in both AMPK and Akt activation, which was accompanied by a remarkable increase in both SIRT3 and SIRT5 expression as compared to the control (Figure 4F-I). Representative western blot images are presented in Figure 4J.

**Figure 4.**
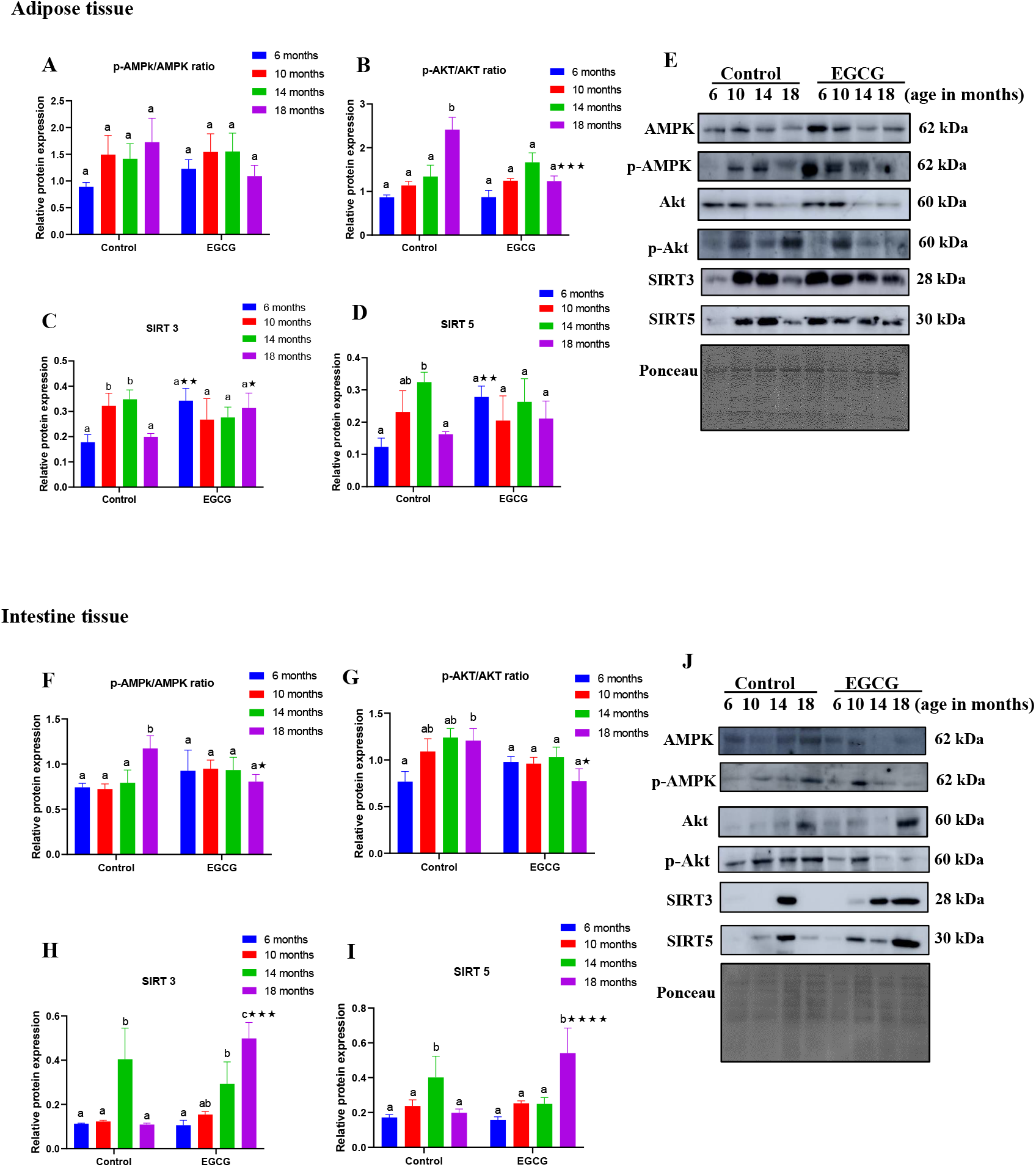
Effect of EGCG consumption on activation of nutrient sensing pathways in various tissues. Animals were divided into 4 control groups and 4 EGCG fed groups, and one group each from control and EGCG groups was sacrificed after every 4 months of feeding till 18 months of animal age. Relative protein expression in adipose tissue **(A)** p-AMPK/AMPK ratio **(B)** p-Akt/Akt ratio **(C)** SIRT3 **(D)** SIRT5 **(E)** Representative western blot images. Relative protein expression in intestine tissue **(F)** p-AMPK/AMPK ratio **(G)** p-Akt/Akt ratio **(H)** SIRT3 **(I)** SIRT5 **(J)** Representative western blot images. Data are presented as means ± SD (n=3) and tested using a 2-way ANOVA followed by Sidak post hoc test. Means that do not share a common letter within a group indicate statistical difference at *P* < 0.05. *represents statistical difference between control and EGCG treated groups at *P* < 0.05.

### 3.5 EGCG administration improves autophagic response in aging animals

Autophagic response was evaluated in terms of LC3II and Atg5 expression in the adipose and intestine tissue. An age-related non-significant decrease in the LC3II expression was evident both in adipose and intestinal tissue, while no discrete trend could be observed for Atg5 expression in the control animals (Figure 5A-F). On the other hand, EGCG consumption significantly enhanced the expression of both LC3II and Atg5 in the intestinal tissue, while only LC3II expression was significantly increased in the adipose tissue on account of EGCG administration (Figure 5A-F).

**Figure 5.**
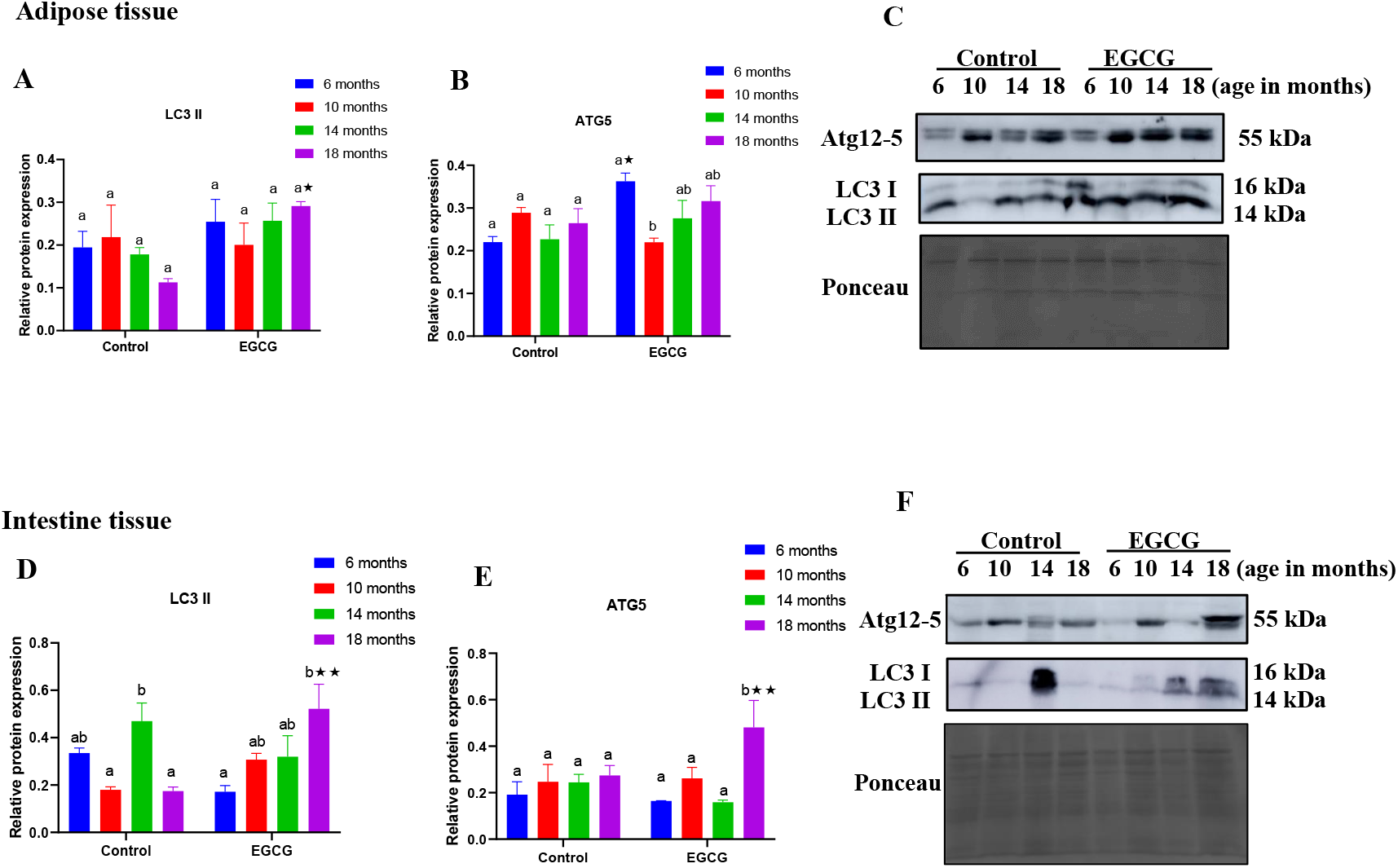
Influence of EGCG consumption on autophagy activation in various tissues. Animals were divided into 4 control groups and 4 EGCG fed groups, and one group each from control and EGCG groups was sacrificed after every 4 months of feeding till 18 months of animal age. Relative protein expression in adipose tissue **(A)** LC3II **(B)** Atg5 **(C)** Representative western blot images. Relative protein expression in intestine tissue **(D)** LC3II **(E)** Atg5 **(F)** Representative western blot images. Data are presented as means ± SD (n=3) and tested using a 2-way ANOVA followed by Sidak post hoc test. Means that do not share a common letter within a group indicate statistical difference at *P* < 0.05. *represents statistical difference between control and EGCG treated groups at *P* < 0.05.

### 3.6 EGCG consumption modulates systemic inflamm-aging and immune cell activation

To further gain insights into healthspan modulating effects of EGCG, several aspects of the immune system were concomitantly analyzed. A robust and significant increase in the circulatory levels of IL-1β and TNF-α was observed in the plasma of 18 months old control group animals as compared to younger animals suggesting prevalent inflamm-aging (Figure 6A,B). Remarkably, this apparent effect appeared to be slightly yet significantly attenuated in EGCG fed animals in the same age group suggesting systemic attenuation of inflammatory damage (Figure 6A,B). Markers of cell senescence were assessed in peritoneal macrophages, and it was observed that although p16^Ink4a^ expression markedly increased in 18 months old animals, however, no such effect was apparent for p53/p21 or β-gal expression (Figure 6C-F). EGCG treatment could not significantly alter either p16^Ink4a^ or β-gal expression although some degree of decrease in p21 expression was evident (Figure 6C-F). Further, no distinct pattern of age-associated changes in the expression of pro-inflammatory genes (TNF-α/IL-6) could be observed in macrophages, while EGCG appeared to slightly decrease TNF-α expression with advancing age (Figure 6 G,H). Immunophenotyping in splenic lymphocytes revealed that total CD3+ cells significantly increased with age in 14 months old animals as compared to younger animals after which a decline in cell numbers was observed (Figure 7A). However, EGCG group animals showed a similar CD3+ T cells profile regardless of age (Figure 7A). The ratio of CD3+CD4+/CD3+CD8+ cells gradually increased with age and was significantly prominent in 18 months old animals while no significant effect of EGCG treatment could be observed (Figure 7B). A remarkable age-associated decrease in NK cells was observed which also could not be significantly altered by EGCG treatment (Figure 7C-K). However, further analyses of CD3+ cells showed that EGCG supplementation enhanced the surface expression of regulatory T cell molecule (CD69) in 18 months old animals as compared to corresponding animals of the control group (Figure 8A-E).

**Figure 6.**
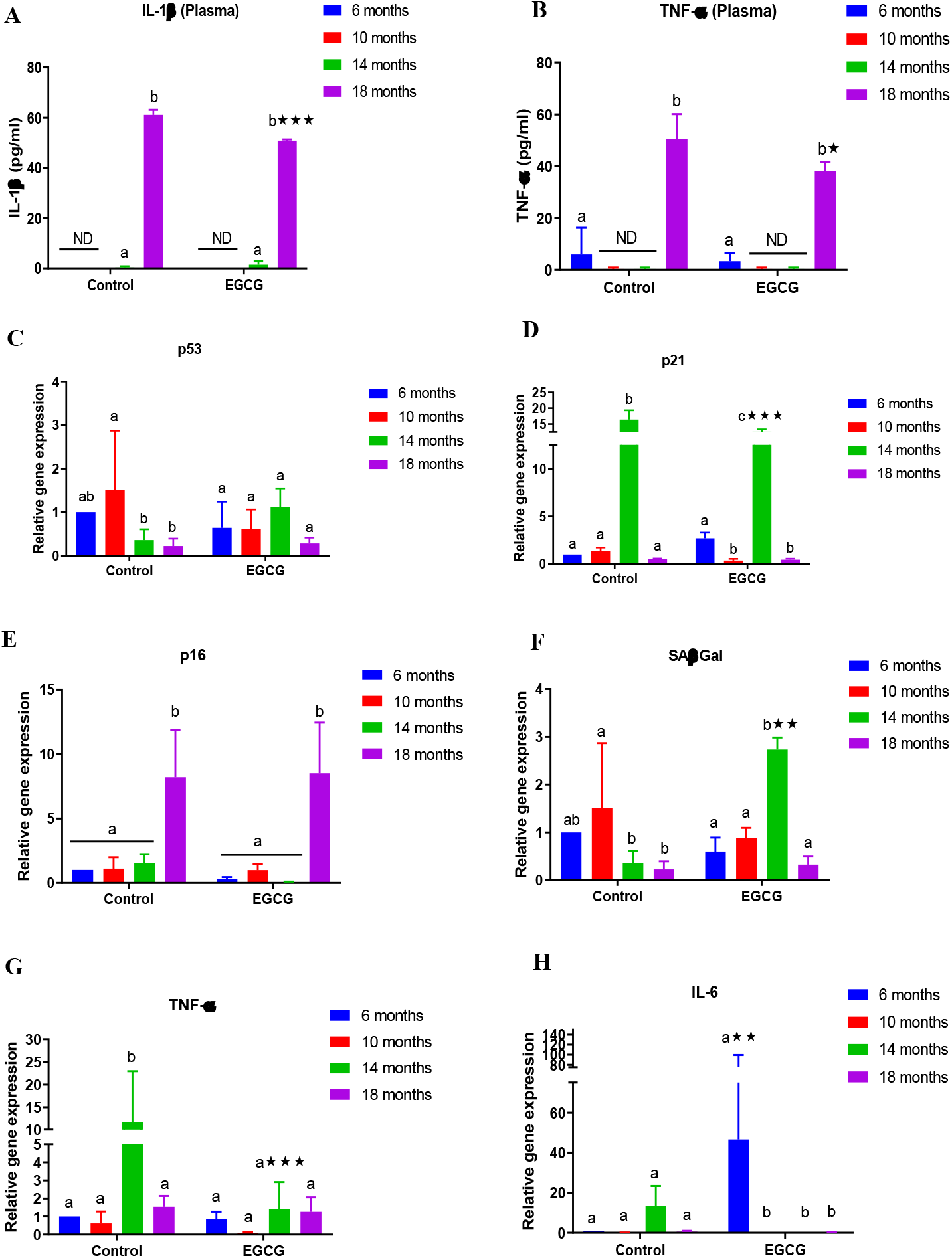
Effect of EGCG consumption on innate immune functions. Animals were divided into 4 control groups and 4 EGCG fed groups, and one group each from control and EGCG groups was sacrificed after every 4 months of feeding till 18 months of animal age. Plasma levels of **(A)** IL-1β **(B)** TNF-α. Relative mRNA expression in peritoneal macrophages of **(C)** p53 **(D)** p21 **(E)** p16 **(F)** SA-β-gal **(G)** TNF-α **(H)** IL-6. Data are presented as means ± SD (n=5) and tested using a 2-way ANOVA followed by Sidak post hoc test. Means that do not share a common letter within a group indicate statistical difference at *P* < 0.05. *represents statistical difference between control and EGCG treated groups at *P* < 0.05.

**Figure 7.**
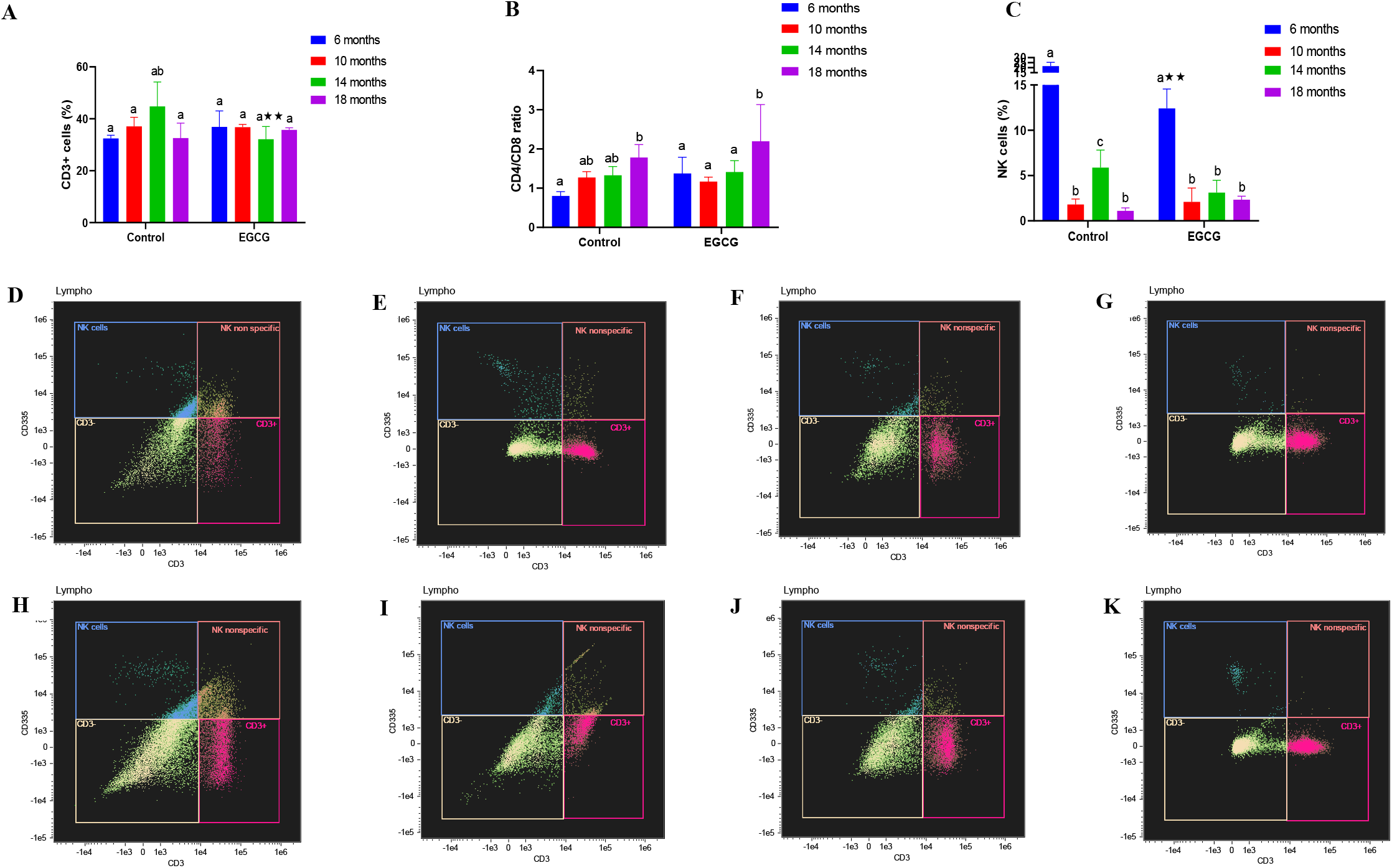
Effect of EGCG consumption on splenic T cell proliferation and activation. Animals were divided into 4 control groups and 4 EGCG fed groups, and one group each from control and EGCG groups was sacrificed after every 4 months of feeding till 18 months of animal age. Abundance of **(A)** CD3+ cells **(B)** CD4/CD8 ratio **(C)** NK cells. Representative flow cytometry plots of NK cells in: Control **(D)** 6 months **(E)** 10 months **(F)** 14 months **(G)** 18 months, and in EGCG: **(H)** 6 months **(I)** 10 months **(J)** 14 months **(K)** 18 months. Data are presented as means ± SD (n=6) and tested using a 2-way ANOVA followed by Sidak post hoc test. Means that do not share a common letter within a group indicate statistical difference at *P* < 0.05. *represents statistical difference between control and EGCG treated groups at *P* < 0.05.

**Figure 8.**
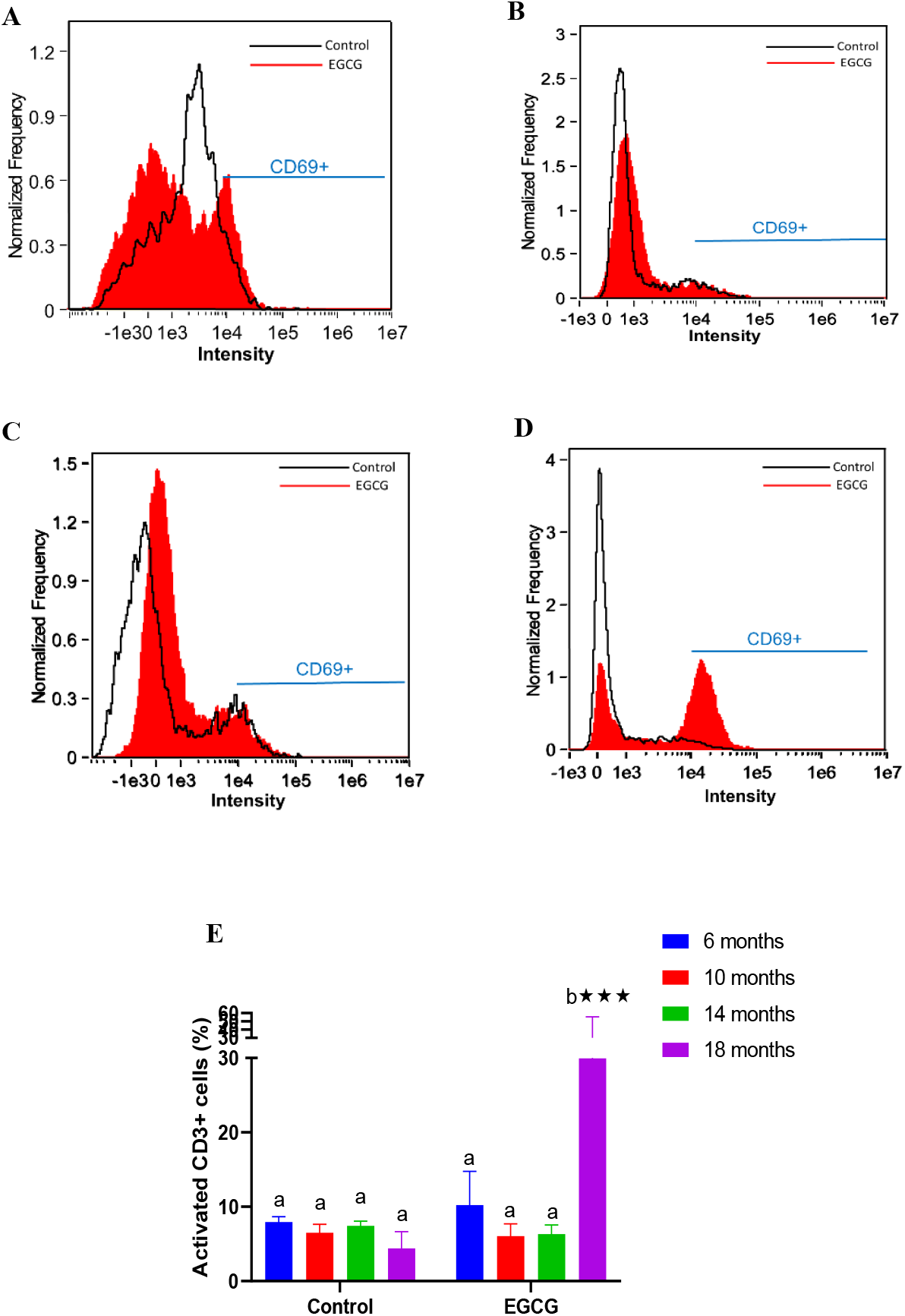
Effect of EGCG consumption on CD69 expression in splenic lymphocytes. Animals were divided into 4 control groups and 4 EGCG fed groups, and one group each from control and EGCG groups was sacrificed after every 4 months of feeding till 18 months of animal age. Representative images of CD69 expression as determined by flow cytometry at age **(A)** 6 months **(B)** 10 months **(C)** 14 months **(D)** 18 months. **(E)** Percentage cells expressing CD69. Data are presented as means ± SD (n=6) and tested using a 2-way ANOVA followed by Sidak post hoc test. Means that do not share a common letter within a group indicate statistical difference at *P* < 0.05. *represents statistical difference between control and EGCG treated groups at *P* < 0.05.

### 3.7 EGCG consumption modulates age-associated changes in the gut microflora

Rarefaction curve analyses of alpha diversity revealed age-associated gradual decline in the richness of microbial species in the control group which was starkly ostensible while comparing 6 months old animals to 18 months old animals (Figure 9A). On the other hand, EGCG treated animals appeared to preserve microbial diversity especially in the 18 months old group as compared to control (Figure 9A). In terms of the beta diversity, presented as PCA plot, 18 months aged control animals showed greater dissimilarity as compared to EGCG fed animals of the same age (Figure 9B). Analyses of microbiome changes at the genus levels showed that the abundance of pathogenic or opportunistic pathogenic genera such as *Clostridium, Staphylococcus, Streptococcus* significantly increased with age in the control groups which appeared to be significantly suppressed in EGCG fed animals, although no effect of EGCG on the relative abundance of commensal microbial genus *Lactobacillus* could be observed (Figure 9C-H).

**Figure 9:**
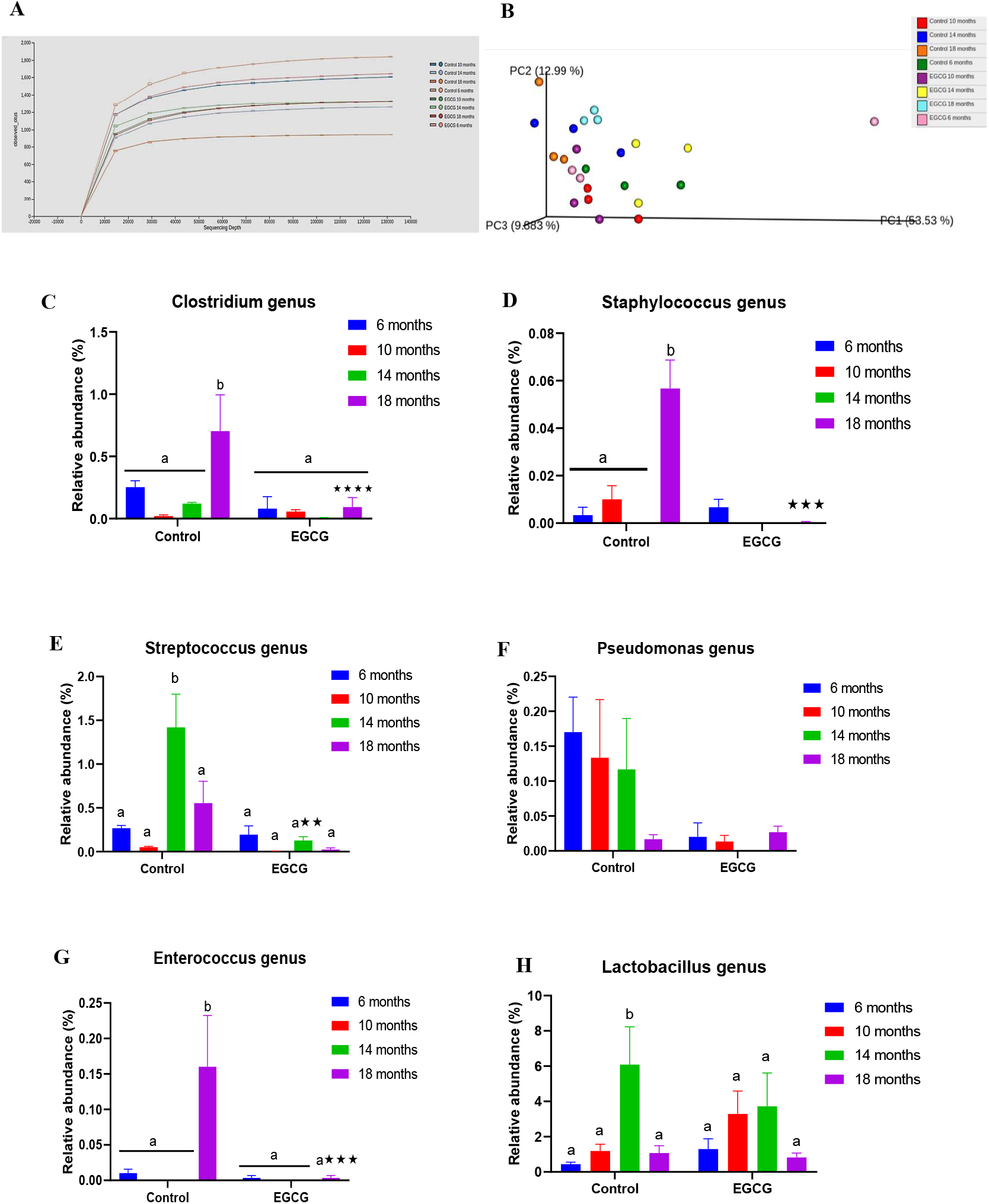
Metagenomic analyses of the gut microbiome. **(A)** Rarefaction curve plot of alpha diversity **(B)** PCA plot of beta diversity. Relative microbial abundance of **(C)** *Clostridium* genus **(D)** *Staphylococcus* genus **(E)** *Streptococcus* genus **(F)** *Pseudomonas* genus **(G)** *Enterococcus* genus **(H)** *Lactobacillus* genus. Data are presented as means ± SD (n=3) and tested using a 2-way ANOVA followed by Sidak post hoc test. Means that do not share a common letter within a group indicate statistical difference at *P* < 0.05. *represents statistical difference between control and EGCG treated groups at *P* < 0.05.

## 4. Discussion

Recent observations suggest that both mitotic and post-mitotic cells can enter a state of cellular senescence, although implications of post-mitotic cellular senescence (PoMiCS) on aging and health are ill-defined (Sapieha and Mallette, 2018). Besides, most of our knowledge regarding PoMiCS is based on studies in neurons and skeletal muscle fibers, while such investigation in post-mitotic adipose tissue in vivo is completely lacking as recently emphasized by von Zglinicki et al (2020). In the present study, an age-associated chronological profile of major mitotic and post-mitotic tissues was undertaken to truly ascertain the extent and depth of prevalent cellular senescence. We observed that the rate of tissue DNA damage, induction of cellular senescence and SASP as well as the modulation of nutrient signaling and autophagic pathways did not appear linear along lifespan but starkly accelerated in tissues of 14-18 months old animals. The prevalent DNA damage in 14-18 months old animals augment the notion that age-related redox imbalance and/or replicative senescence-induced ROS generation can significantly contribute to the initiation of cellular senescence. It was also clearly noticeable that adipose tissue is a highly vulnerable site of spontaneous age-associated DNA damage, development of cellular senescence as well as SASP followed by the intestinal tissue while liver appeared to be least affected. As far as we know, this is the first report to suggest that post-mitotic adipose tissue strongly represents characteristic features of cellular senescence in vivo which enhances our understanding of the global physiological impact of cellular senescence. This is also of particular interest since our study did not use any obesity/high fat diet model that may interfere with normal adipose functions, and yet adipose tissue presented with maximum induction of p53/p21 dependent senescence pathway and SASP. This also provides a new perspective to a previous report by Minamino et al (2009) which showed that excessive calorie intake increased oxidative stress in the adipose tissue of mice with type 2 diabetes–like disease and promoted senescence-associated changes, including increased expression of p53 and production of proinflammatory cytokines. It is thus evident that the pathological role of adipose tissue, in the context of cellular senescence, is independent of its physiological abundance or related underlying disease condition. In addition, it has already been observed that senescent preadipocytes can directly influence the aging phenotype in immune cells such as macrophages (Kumar et al 2020), and that elimination of senescent cells can alleviate obesity-induced type II diabetes suggesting multifaceted harmful implications of senescent adipose tissue (Palmer et al 2019). Although age-associated changes in the morphology and functional inefficacies of intestinal epithelium are well known; reports addressing in vivo cellular senescence in intestinal tissue are rare. A previous study by Moorefield et al (2017) showed that mRNA expression of cell cycle inhibitor p53 and marker of oxidative stress (Nrf2) are upregulated in intestinal epithelial stem cells of aged mice. A recent study in mice exposed to radiation revealed the development of premature senescence and SASP in intestinal stem cells (Kumar et al 2019). However, our observations indicate that intestinal tissue is also disposed to age-induced activation of cellular senescence and SASP which may explain the known functional deterioration of intestinal epithelium with gradual aging. On the contrary, no robust activation of cellular senescence or SASP could be established in the liver despite an evidence of p53 upregulation and slight inflammatory activation. These variations in different cell types could be attributed to different rates of tissue renewal and/or susceptibility to different triggers of senescence. Liver is a robust organ and its functions are largely preserved with age despite 20-40% shrinkage of liver volume, reduction in cytochrome c activity and a gradual increase in serum bilirubin with age (Wynne et al 1989; Boland et al 2014). However, age-related susceptibility of liver to fibrosis and chronic disorders including NAFLD has been recognized which is attributed to reduced regenerative capacity and immune dysfunctions (Allaire and Gilgenkrantz, 2020). Besides, being a major metabolic organ and detoxification center, a sophisticated antioxidant defense system protects the liver from redox imbalance which might contribute to its prolonged protection from oxi-inflammatory stressors, and therefore delay the onset of cellular senescence/SASP as observed in the present study. In fact, in vivo cellular senescence in liver has been considered more relevant in the pathogenesis of various liver disorders particularly in the case of hepatic steatosis (Ogrodnik et al 2017; Papatheodoridi et al 2020). Given the strong upregulation of inflammatory factors courtesy of SASP in the adipose and intestine tissues, we also speculate that these senescent tissues may significantly contribute to the systemic prevalence of pro-inflammatory factors (inflamm-aging) as we observed in this study.

Daily consumption of EGCG in the present study dramatically improved several aspects of healthspan and lifespan in mice. EGCG consumption significantly decreased the average risk of death in animals and thus enhanced lifespan which was remarkably similar to a previous study on rats (Niu et al 2013). In the context of cellular senescence, EGCG consumption strongly downregulated the DDR, induction of cell cycle inhibitory and SASP pathways albeit with varying degree in the investigated tissues suggesting a robust systemic effect. We and others have previously shown that EGCG can attenuate premature senescence in preadipocytes and fibroblasts in vitro by downregulating p53/p21 pathway as well as SASP (Han et al 2012; Kumar et al 2019). However, to our knowledge, this is the first in vivo study to assess the effects of EGCG consumption on different aspects of cellular senescence, and our results suggest that EGCG enhances lifespan by attenuating the onset of cellular senescence and associated damage in multiple tissues. We also observed a strong reduction in the activation of SASP related inflammatory transcription factors (p38MAPK/ NF-κB) and effector molecules (cox-2) in tissues of EGCG treated mice. A concomitant decrease in circulatory inflammatory factors in plasma further provided strong evidence of anti-inflamm-aging and anti-SASP effects of EGCG. SASP is a particularly detrimental aspect of cellular senescence as it can induce a bystander effect and promote a chronic pro-tumorigenic and pro-inflammatory environment in nearby healthier cells (Coppé et al 2010). Multiple stress signals can activate p38MAPK which is recognized as a necessary and sufficient activator for a stable SASP expression in senescent cells (Freund et al 2011). p38MAPK can directly regulate NF-κB activity and together, these transcription factors present a robust SASP activating mechanism. EGCG has been shown to modulate p38MAPK and NF-κB expression in various experiment settings, and our results corroborate the anti-inflammatory attributes of EGCG in the context of SASP thereby providing a deeper understanding of its pharmacological effects during aging. Together, it is reasonable to assert that chronic EGCG consumption has the potential to counter both cellular senescence and SASP, and thus EGCG is a unique candidate for developing cellular senescence based anti-aging strategies.

We next assessed nutrient sensing pathway mediators to understand the mechanism(s) of observed effects of EGCG. AMPK and Akt activation were apparent in senescent tissues indicative of prevalent metabolic stress. Senescent cells display altered metabolic features characterized by increased ADP:ATP and AMP:ATP ratios which activate the master regulator AMPK resulting in a pro-glycolytic but less energetic state (Zwerschke et al. 2003). The activated AMPK can also induce the expression of p53/p21 cell cycle inhibitors by at least two known mechanisms that may initiate the cellular senescence program (Wiley and Campisi 2016). In addition, AMPK has also been shown to actively phosphorylate Akt under various cellular stressors, and persistent activation of Akt suppresses FOXO3a transcription factor and increases levels of intracellular ROS which subsequently initiate p53/p21 dependent growth arrest (Miyauchi et al 2004). It is thus not surprising that inhibition of Akt/mTOR is considered amongst the best-known interventions to extend lifespan. As such, in our previous in vitro study, we reported that EGCG treatment can attenuate premature cellular senescence in preadipocytes by inhibiting Akt activation which is corroborated in the present study (Kumar et al 2019). Nutrient sensor sirtuins have also been associated with increased lifespan, and mitochondrial sirtuins SIRT3 and SIRT5 are emerging as critical regulators of mitochondrial deficits associated with aging (Shih and Donmez, 2013). In the present study, SIRT3 activity was strongly upregulated in tissues of EGCG fed animals which can induce mitochondrial enzymes involved in the electron transport chain, antioxidant defenses and energy metabolism thereby augmenting mitochondrial functions and redox homeostasis with age. A recent study by Lilja et al (2020) has also shown that EGCG application can reduce premature senescence and SASP in preadipocytes by activating SIRT3 expression. Similar to SIRT3, SIRT5 is emerging as an important regulator of mitochondrial respiration and redox balance, although its role in senescence and aging is not well defined. SIRT5 overexpression has been shown to promote desuccinylation and activate Cu/ZnSOD to eliminate ROS (Lin et al 2013), whereas a lack of SIRT5 results in increased ROS production and susceptibility of affected cells to oxidative damage (Zhou et al 2016). It is thus reasonable to suggest that the observed anti-cellular senescence attributes of EGCG in the present study could be mediated by maintenance of robust mitochondrial functions, and thus the redox homeostasis owing to the observed strong upregulation of SIRT3/5. It also needs to be emphasized here that activation of sirtuins is one of the key mechanisms of calorie restriction-induced increase in the lifespan, and our results suggest that EGCG could act as a potential calorie restriction mimetic. In addition to accumulation of senescent cells, impaired mitochondrial functions, redox imbalance and alterations in nutrient sensing pathways; cellular autophagic response is also influenced with age. It is thus not surprising that activation of basal autophagy has been reported to enhance lifespan and improve age-related pathologies in vital organs (Fernandez et al 2018). Our results showed robust activation of autophagic flux as indicated by enhanced LC3II and ATG5 expression especially in the tissues of 18 months old EGCG fed animals. Previous studies report that EGCG can induce autophagy by promoting the synthesis of autophagosomal protein LC3II (Zhao et al 2017), autophagy-dependent cell survival (Holczer et al 2018) and in fact, the anti-inflammatory potential of EGCG has also been linked to enhanced autophagic response (Li et al 2011). Taken together, it can be concluded that EGCG consumption modulates age-related impairment in multiple nutrient sensing pathways and autophagic response to ultimately induce metabolic and oxi-inflammatory homeostasis which attenuate the development and progression of cellular senescence/SASP and thus enhance organismal longevity. Unlike other cells, characterization of cellular senescence and its biological relevance in the immune cells has been less explored. In particular, senescent macrophage identification in vivo has been disputable as presence of the most often used senescence markers have been described as an intrinsic, reversible property of macrophages (Burton and Stolzing, 2018). We sought to determine the extent of cellular senescence in peritoneal macrophages by analyzing mRNA expression of cell cycle inhibitors and mediators of inflammation, but unlike other tissues, we could not observe any discernible pattern. Despite evidence of p16^Ink4a^ activation with advancing age in macrophages, our results implicate further deeper studies to understand macrophage cellular senescence vis-à-vis immunosenescence. On the other hand, an age-associated increase in the CD4+/CD8+ T cells ratio, and a gradual decline in NK cell numbers were observed, which did not appear to be influenced by EGCG. However, EGCG supplementation enhanced the expression of regulatory molecule CD69 on T cells in 18 months old animals which has also been reported for different natural molecules (Holderness etal 2011; Ramstead et al 2015; Tumová et al 2017). CD69 is considered a marker of early T cell activation although evidence of its more dynamic and regulatory effects is also emerging. CD69 expression can directly impact the migration of immune cells (Shiow et al 2006), and CD69 deficient mice show exacerbated pro-inflammatory immune response (Radulovic et al 2013). Further, a higher pro-inflammatory environment has been recognized in T cells in elderly (Zanni et al 2003) and therefore, the apparent overexpression CD69 in EGCG treated T cells could be relevant in attenuating the exaggerated pro-inflammatory environment in immune cells. Metagenomic microbiota analyses showed that EGCG fed animals preserved the age-associated loss in the diversity of microorganisms, while simultaneously suppressing a number of pathogenic/opportunistic pathogenic microbes. Gut microbiota dysbiosis has been directly linked to the induction of cellular senescence and diseases (Yoshimoto et al 2013; Tan et al 2019), and potential modulators of the gut microbiome may have anti-cellular senescence attributes (Sharma and Padwad, 2020c). In addition, polyphenols are being advocated as promising alternative to oligosaccharide-based prebiotics, and previous studies have shown that polyphenols can influence probiotic growth either directly or by suppressing the abundance of pathogenic bacteria (Sharma and Padwad, 2020d). The present study suggests some degree of correlation between the observed effects of EGCG in tissues and the gut microbiota modulation, although further work is required to establish a causative relationship.

## 5. Conclusion

In summary, our study presents compelling evidence of prevalent cellular senescence and SASP in post-mitotic adipose tissue, and also report that EGCG consumption can counter several deleterious aspects of cellular senescence, SASP and inflamm-aging. These attributes of EGCG could also be linked to improvement in immune functions and maintenance of gut dysbiosis, although deeper analyses are warranted for a substantial correlation. Nonetheless, it is evident that EGCG consumption has several unique and multifaceted cellular senescence mitigatory attributes which may improve both longevity and healthspan.

## Abbreviations

SC: Senescent cells
SASP: Senescence associated secretory phenotype
EGCG: Epigallocatechin gallate
AMPK: 5’-Adenosine monophosphate-activated protein kinase
Cox-2: Cyclooxygenase-2

## Funding

This work was supported by grant from the Department of Science and Technology, Government of India under the INSPIRE Faculty scheme (IFA17-LSPA79).

## Declaration of competing interest

The authors declare that they have no competing interests.

## Acknowledgements

Authors are grateful to the director of Council of Scientific & Industrial Research-Institute of Himalayan Bioresource Technology for constant support and encouragement. This work was supported by grant from the Department of Science and Technology, Government of India under the INSPIRE Faculty scheme (IFA17-LSPA79).

## Notes

### Competing Interest Statement

The authors have declared no competing interest.

